# First report of the sexual stage of the flax pathogen *Mycosphaerella linicola* in France and its impact on pasmo epidemiology

**DOI:** 10.1101/2020.06.17.156984

**Authors:** Delphine Paumier, Blandine Bammé, Annette Penaud, Romain Valade, Frédéric Suffert

## Abstract

We performed a three-year field survey in France to characterize the dynamics of sexual reproduction in *Mycosphaerella linicola*, the causal agent of pasmo, during the interepidemic period. Cohorts of fruiting bodies were sampled from linseed straw during the autumn and winter and carefully observed, focusing on pseudothecia, asci and ascospores. A sequence of experimental steps corresponding to Koch’s postulates confirmed in July 2014, for the first time in France and continental Europe, the widespread presence of the sexual stage of *M. linicola* in plant host tissues. The developmental dynamics of pseudothecia on straw, expressed as the change over time in the percentage of mature pseudothecia, was similar in all three years. Pseudothecia appeared in late summer, with peak maturity reached in October. A temporal shift, thought to be due to early autumn rainfall, was highlighted in one of the three years. These observations suggest that sexual reproduction plays a significant role in the epidemiology of pasmo in France. A resurgence of *M. linicola* infections in spring flax is thought to have occurred in recent years, due to the increase in the area under flax. The presence of the sexual stage of this pathogen probably increased the quantitative impact of residues of winter linseed (used for oil) and flax straw (left on the soil for retting and used for fibers) as an interepidemic ‘brown bridge’. This case study highlights how certain parts of a disease cycle, in this case the sexual phase, can become crucial due to changes in production conditions.

## Introduction

*Mycosphaerella linicola* Naumov, anamoph *Septoria linicola* (Speg.) Garassini, is the causal agent of pasmo, a disease affecting both ‘flax’ and ‘linseed’ (*Linum usitatissimum* L.), crops grown for fiber and oil production, respectively. This plant-pathogenic ascomycete affects production in many flax-growing areas around the world, including Europe. Pasmo was first detected in Argentina in 1909 (Spegazzini, 1911). In Europe, it was found in Yugoslavia in 1936 (Rost, 1937) and in Ireland in 1946 (Loughnane et al., 1946). More recently, this disease has caused significant yield losses in South Dakota (Ferguson et al., 1987) and in England, on winter linseed (Perryman & Fitt, 1999).

Pasmo usually starts on the lower leaves of young plants. During the growing season of the plant, *M. linicola* is propagated clonally by splash-dispersed asexual pycnidiospores (filiform 1-3 septate, straight or slightly curved conidia; Sivaneson & Holliday, 1981), leading to an upward progression of the disease on stem and leaves. Pasmo affects all the aboveground parts of the flax plant, causing leaf spots, a loss of flowers and capsules, and weakened pedicels (Fergusson et al., 1987). The disease also causes elongated brown lesions, which coalesce to produce mottled bands that encircle the stem (Colhoun & Muskett, 1943; Perryman et al., 2009). This ‘girdling’ effect and the premature death of the plant has an impact on yield and fiber quality, and may be mistaken for premature ripening (Sackston & Carson, 1951; Perryman et al., 2000).

*M. linicola* has been reported to be seed-borne and to survive on crop residues, like many other pathogenic ascomycetes. The sexual stage was initially reported to be absent in North Dakota (Brentzel, 1926), but several plant pathologists identified structures found on flax stubble as sexual fructifications, in Argentina (Wollenweber, 1938) and Germany (Kruger, 1941). Sackston (1949) identified structures that he believed to be ‘pseudothecia’ in Manitoba (Canada), but he was unable to complete Koch’s postulates. Despite partial evidence for a sexual stage being repeatedly reported, this pathogen has long been thought to survive the interepidemic period on infected flax straw as a saprotroph (mycelium and old pycnidia; Gillis, 2009), with dispersal purely by rain splash or insects during the growing season (Christenson, 1952; Sackston, 1970), and long-distance carriage only in or on seeds. Sanderson (1963) was the first to identify and describe asci and ascospores from wild flax (*Linorum marginale*) in New Zealand, but, unfortunately, was unable to establish cultures. Pseudothecia resemble black spherical dots, 75-120 μm across, whereas asci are oblong, bitunicate, eight-spored and 30-50 × 8-9 μm in size, and ascospores are fusiform, hyaline two-celled, constricted at the septum, and 13-17 × 2.5-4.0 μm in size (Sivaneson & Holliday, 1981; Vakhrusheva, 1986). It was not until Perryman et al. (2009) collected air-borne ascospores thought to be those of *M. linicola* that a significant role of the sexual stage in the pasmo epidemiology could be affirmed. These authors proposed a pathogen life cycle including a sexual phase during the winter, in which the pathogen survives as pseudothecia, although these fruiting bodies were not observed in their study. An important role of the sexual stage and the recombination resulting from it is consistent with the high levels of genetic diversity demonstrated for two populations from Manitoba (Grant, 2008).

Pasmo incidence and severity have increased over the last two decades in France (no attack reported in French trials in 1988 and 1990; Fitt et al., 1991), in parallel with the increase in the area under *Linum usitatissimum* L., particularly in the form of flax for linen production (50,000 ha in the 1990s vs. 120,000 ha in 2020; data CIPALIN). During this period, France has become the world’s largest producer of scutched flax fibers (580,000 t in 2017, i.e. about 75% of the world’s production; FAOSTAT). Nevertheless, the sexual stage of *M. linicola* – which probably plays a key role in the epidemiology of this disease – has never before been described in France, or elsewhere in continental Europe.

In this study, we aimed to characterize the sexual reproduction dynamics of *M. linicola* on flax straw during the interepidemic period, to assess its role in the early stages of pasmo epidemics in French production conditions.

## Materials and methods

### Description of the different forms of M. linicola during the intra- and interepidemic periods

Pasmo symptoms and both the asexual and sexual stages of *M. linicola* were observed on flax stems, leaves and straw collected from two fields in 2015-16 at the INRAE-Terre Inovia experimental station (Thiverval-Grignon, France). Fruiting bodies were collected with a needle, crushed and examined under a microscope, after mounting in methylene blue solution. Pycnidia and pycnidiospores (**Figure 1A-B**) were obtained from typical lesions on living flax leaves. Pseudothecia, asci and ascospores were identified on pieces of flax straw left to dry outside (**Figure 1C-G**).

**Figure 1.**
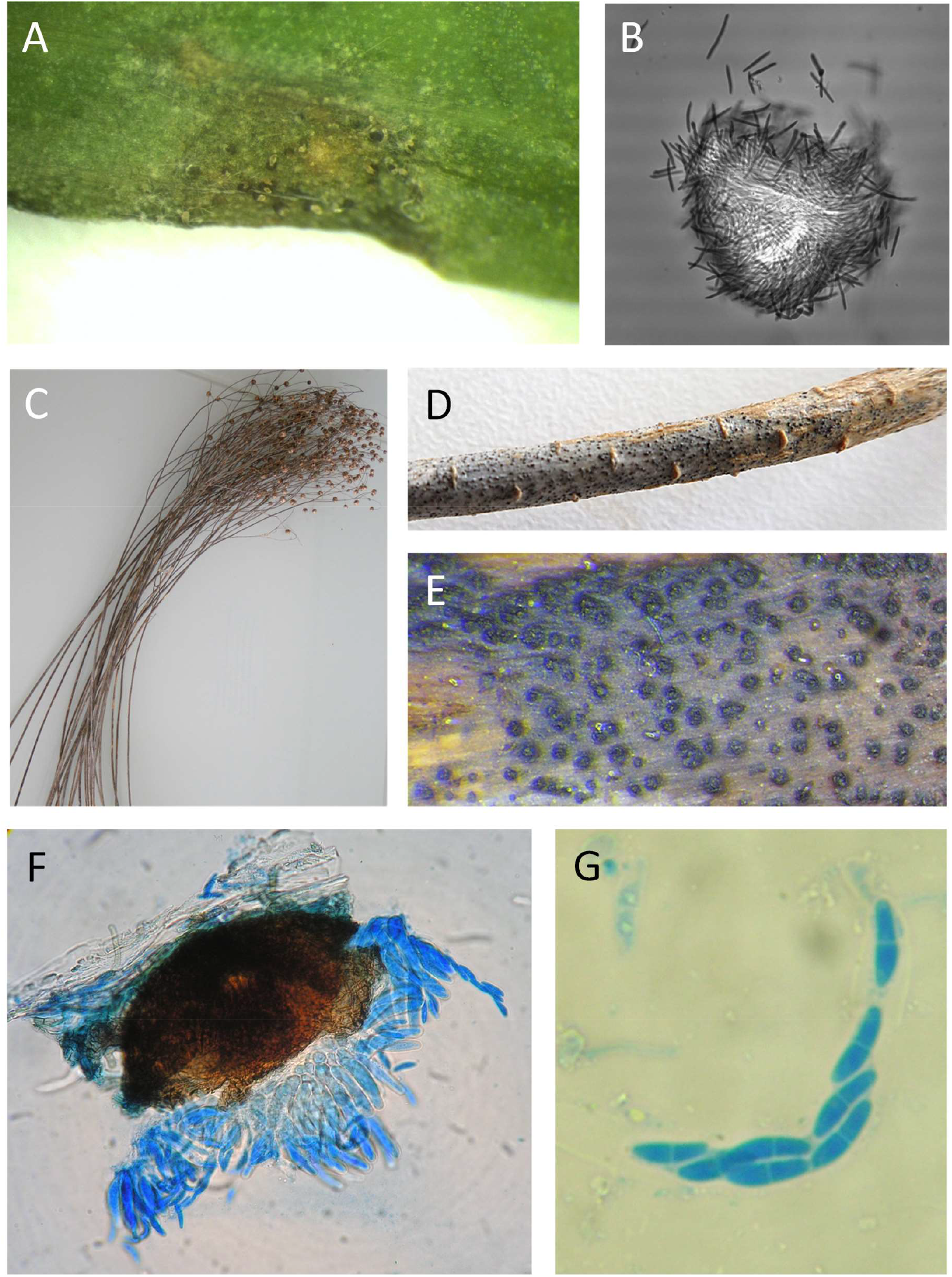
A. Early septoria lesion on a flax leaf with pycnidia exuding *Mycosphaerella linicola* cirrhi. B. Pycnidium, showing a cluster of several hundred pycnidiospores (×10). C. Flax (*Linum usitatissimum*) straws collected from a field during retting. D, E. Pseudothecia (black points) on flax stem residues. F. Mature pseudothecium (brown) and asci (×400). G. Ascus containing 8 ascospores (blue).

### Sequence of experimental steps demonstrating the presence of the sexual form

A sequence of experimental steps corresponding to classical Koch’s postulates was developed to confirm that the observed fruiting bodies did indeed correspond to the sexual stage of *M. linicola* and could potentially play a role in pasmo development (**Figure 2**). Flax stem residues bearing putative *M. linicola* pseudothecia were identified (➀). We used a technique for ascospore collection derived from that developed by Suffert & Sache (2011) to study the sexual stage of *Zymoseptoria tritici* on wheat residues to isolate them. Dry fragments of flax straw were spread on wet paper in a moist box (24 × 36 cm). Petri dishes containing PDA medium (potato dextrose agar, 39 g L^-1^) were placed upside-down above the fragments (➁). The box was placed at 18°C in the dark for 12 h. The Petri dishes were closed and incubated in the same conditions. Three days later, white yeast-like colonies similar to those obtained by Wollenweber (1938) from *S. linicola* pycnidiospores appeared (➂). A conidial suspension was prepared by flooding the surface of a 3-day-old culture of a single-spore colonies obtained after one round of single-spore isolation with sterile water and scraping the surface of the agar with a glass rod (➃). Leaves of a flax plant grown in a greenhouse were inoculated by applying this conidial suspension to the adaxial surface, with a paintbrush (➄). The plant was enclosed in a transparent polyethylene bag containing a small amount of water to maintain humidity levels and, thus, promote infection. Pycnidia and cirrhi, accompanied by symptoms, appeared on the inoculated leaves 10-13 days after inoculation (➅). Cirrhi were collected with a needle (➆) and deposited on PDA medium in a Petri dish for reisolation of the pathogen in its yeast-like form (➇). Clusters of several hundred pycnidiospores released from other cirrhi mounted in methylene blue solution were examined under the microscope. At the same time, putative pseudothecia were observed on flax straw collected in the field and examined in a similar manner (➈).

**Figure 2.**
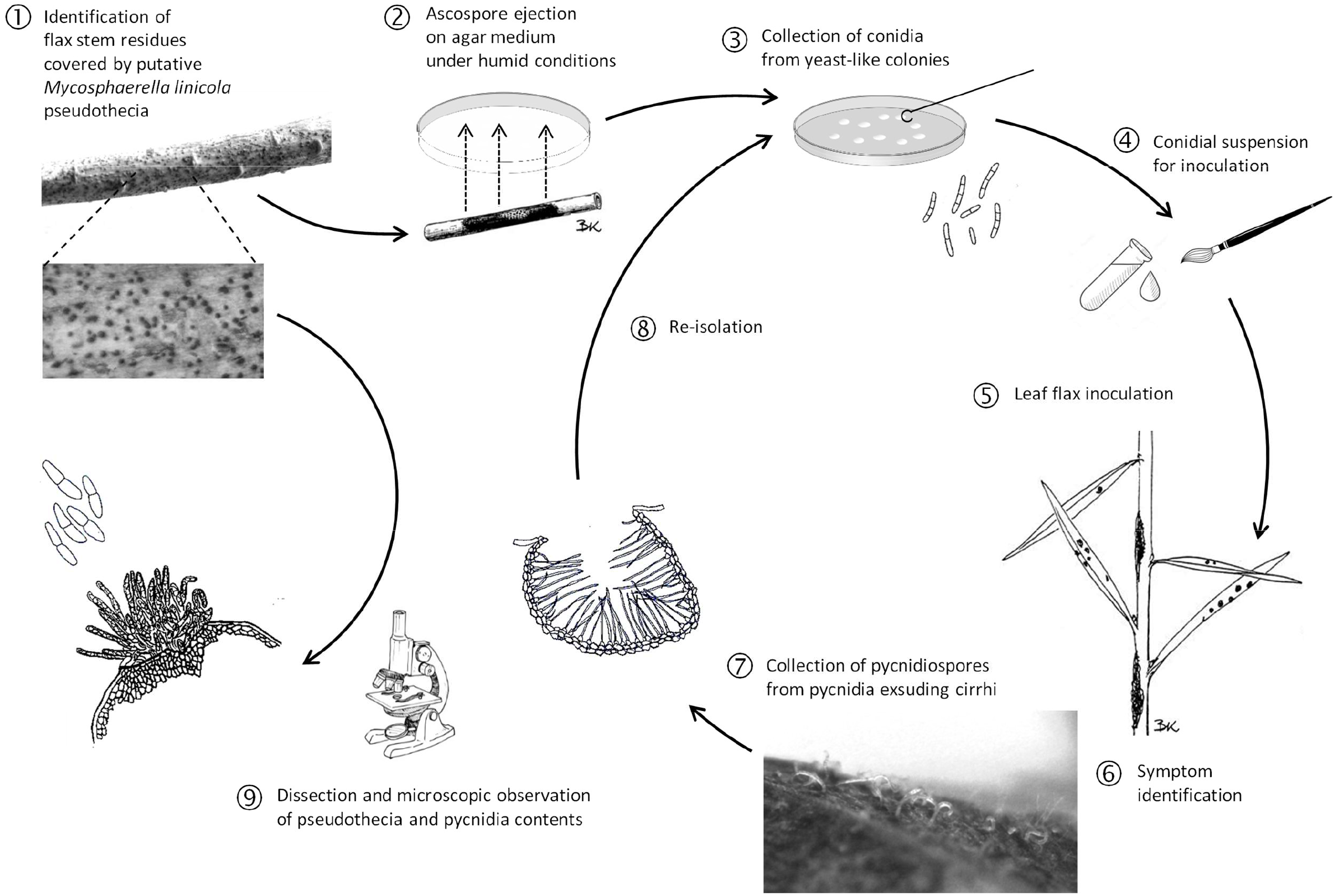
Sequence of experimental steps (Koch’s postulates) demonstrating the relationship between the presence of the sexual form of *Mycosphaerella linicola* on flax stem residues and pasmo symptoms on leaves.

### Dynamics of M. linicola on flax straw during the interepidemic period in French conditions

During three growing seasons (2014-15, 2015-16, 2016-17) straw was collected from flax naturally infected with *M. linicola* in a field planted with a winter linseed varietal mixture (cv. Blizzard, Sideral, Cristalin, and Angora). The straw was collected just after harvest in July and stored outdoors, on grass, at the INRAE-Terre Inovia experimental station. In 2014-15, the straw was moistened before storage, whereas, in 2015-16 and 2016-17, it was left on the ground directly, without treatment. The straw was examined once weekly from 30 October 2014 to 26 March 2015 for the 2014-15 season, from 27 August 2015 to 6 April 2016 for the 2015-16 season, and from 21 July 2016 to 9 March 2017 for the 2016-17 season. An automatic weather station located 200 m from the pile of straw recorded hourly rainfall and air temperature at a height of 2 m. Each week, we observed a sample of 20 straws under a binocular microscope, to check for the presence of *M. linicola* pycnidia and pseudothecia. As soon as these structures appeared, a subsample of five infected straws was incubated in a moist box for 24 h. Five fruiting bodies were selected from each straw (25 in total) and dissected under a binocular microscope according to the protocol developed by Poisson (1997) for *Leptosphaeria maculans*. Fruiting bodies were placed in a drop of water on a glass slide and crushed with two needles to release their contents. A drop of methylene blue solution was added and the slide was covered with a coverslip. The preparation was observed under a microscope (magnification ×400), to distinguish pycnidia from pseudothecia. The *M. linicola* pseudothecia were classified on the basis of developmental stage, taking into account the degree of maturity of asci and ascospores, as proposed by Toscano-Underwood et al. (2003) for *L. maculans* and *L. biglobosa*. Four classes were defined and used: (i) ‘immature’ pseudothecium, with no asci or ascospores; (ii) ‘maturing’ pseudothecium, with differentiated asci but no ascospores; (iii) ‘mature’ pseudothecium, with at least one ascus containing eight differentiated ascospores; (iv) ‘empty’ pseudothecium, from which all the ascospores had been discharged (**Figure Suppl. 1**). A pseudothecium was considered to have reached one of the four developmental classes when the first asci/ascospores in the pseudothecium had reached the stage of development considered.

## Results

*M. linicola* pseudothecia and ascospore-containing asci were observed in July 2014 for the first time in France, and are described here (**Figure 1 and 3**). Koch’ postulates were completed and showed that ascospore-derived strains were able to generate typical pasmo symptoms on flax leaves after inoculation with a paintbrush and exposure to high-moisture conditions (**Figure 2**). Based on weekly counts of *M. linicola* fruiting bodies (pycnidia and pseudothecia) on samples of 20 infected straws, and the dissection of five pieces of straw under a binocular microscope, we were able to determine the dynamics of pseudothecium formation and maturation over the interepidemic period, from August to April, in three successive growing seasons (2014-15, 2015-16, 2016-17) (**Figure 4**). This epidemiological survey provides the first evidence for the widespread presence of the sexual stage of *M. linicola* in France just before the emergence of the winter flax crop. The highest proportion of mature pseudothecia, ranging from 60% to 100%, was recorded in October, regardless of the year considered. The dynamics of pseudothecium maturation were similar in 2014-15 and 2015-16, with a peak in early October, followed by a steady decrease to below 20% after December. The peak was delayed by one month (early November) in 2016-17. The 2014-15 survey did not begin until early November, and we were, therefore, unfortunately, unable to determine when peak pseudothecium maturity occurred. The first symptoms of pasmo were detected on flax seedlings (cotyledons) in the field on 25 November, 2015 (second growing season) and on 14 December, 2016 (third growing season). A similar assessment was performed in the first growing season, but at a later stage, making it impossible to determine whether symptoms occurred as early in the autumn as in subsequent seasons. The identification of *M. linicola* was confirmed by a TaqMan qPCR assay. The intron sequence of the EF1-α gene was amplified using the specific primers ‘SeptoUP’ (5′-TTGCCCCTCCAATTCTGGTG-3′), ‘SeptoLOW’ (5’-ATGTGTTAAAAGTGTTGTGTGC-3’), and the TaqMan probe ‘Sonde_Septo’ (5’-FAM-CGAGAATTTTGGGCTTTTGCGGCTC-BHQ1-3’). The specificity of primers was confirmed with DNA extracted from pure cultures of 50 strains of *M. linicola* and of 24 other fungal species, including the wheat pathogen *Z. tritici* and the flax pathogens *Sclerotinia sclerotiorum, Rhizoctonia* sp., *Verticillium dahliae, Pythium* sp., *Phoma exigua* var. *linicola* and *Kabatiella lini*.

**Figure 3.**
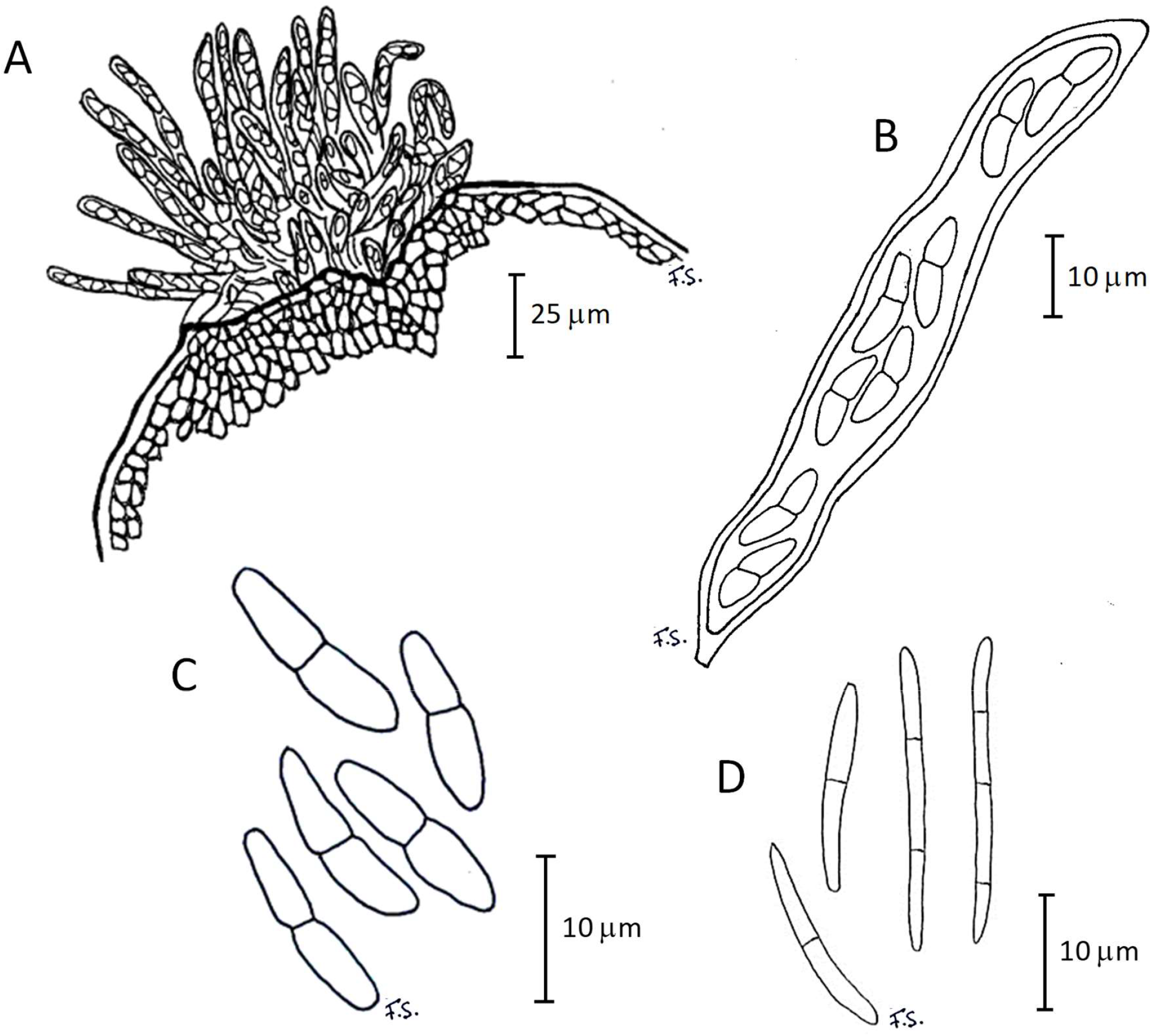
*Mycosphaerella linicola* sexual fructifications. A. Pseudothecium; B Ascus; C. Ascospores; D Pycnidiospores.

**Figure 4.**
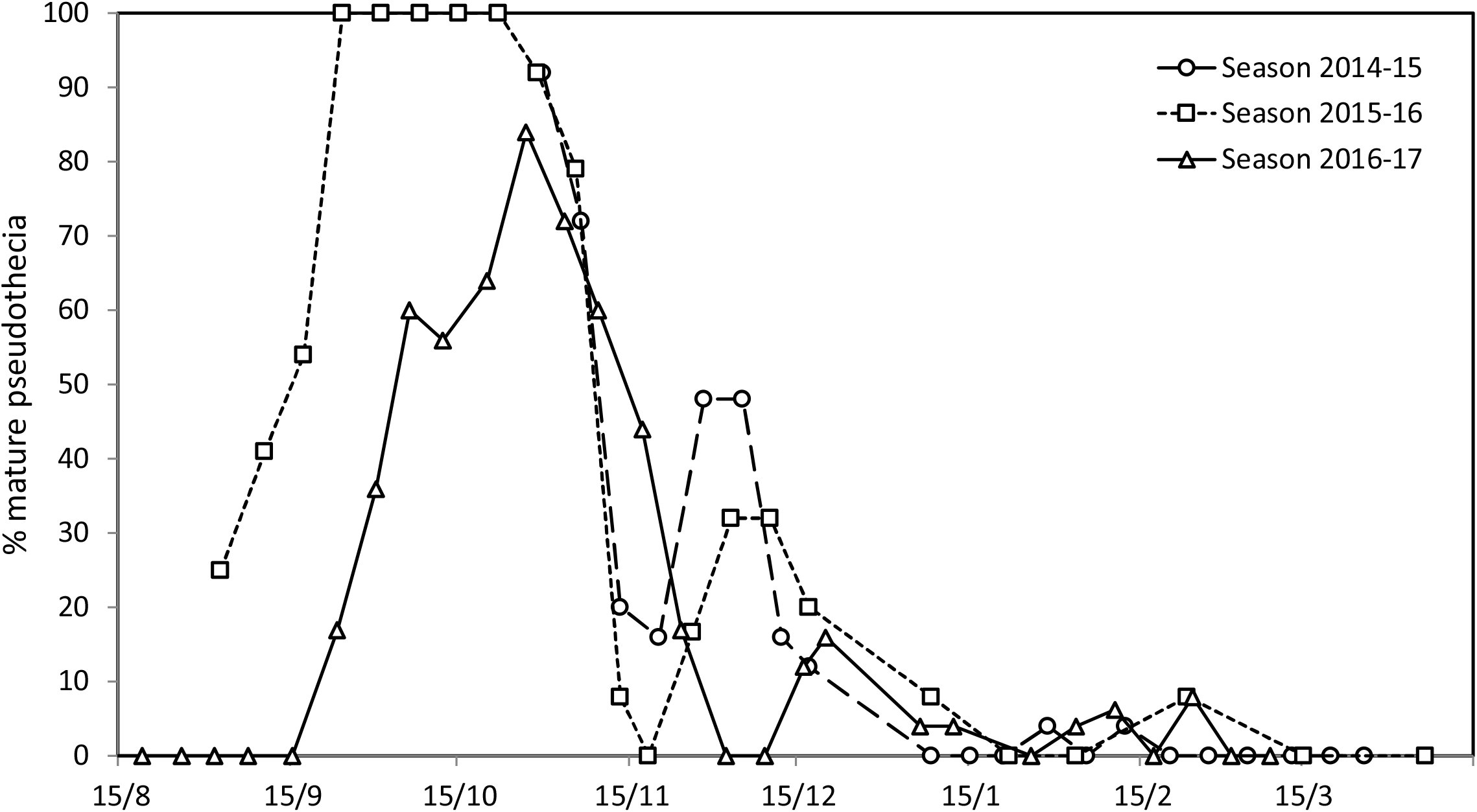
Dynamics of the sexual stage of *Mycosphaerella linicola* on flax straw during the 2014-15, 2015-16 and 2016-17 growing seasons (from July to April), expressed as the change in the percentage of mature pseudothecia over time (see **Figure Suppl. 1**).

The year in which symptoms were detected earliest on emerging plants (late November 2015 vs. mid-December 2016) was also the year in which pseudothecium levels peaked earliest (mid-September 2015 vs. late-October 2016). This suggests that early symptoms may be directly related to the early availability of ascospores to act as a primary inoculum, as demonstrated for other ascomycete pathogens of plants, including *Z. tritici* (Morais et al., 2016), *Pyrenophora tritici-repentis* (Adee & Pfender, 1989) and *L. maculans* (Naseri et al., 2009). However, the observations reported here should be interpreted with caution, because flax seedlings emerged more than three weeks later in the 2016-17 season than in the 2014-15 and 2015-16 seasons, due to a lack of rain after sowing (**Figure Suppl. 2**). Moreover, weather conditions contrasted strongly between the three growing seasons at Thiverval-Grignon. In the period before flax crop emergence in the 2016-17 season, July and August were particularly dry, with a total rainfall of only 32 mm (**Figure Suppl. 2**). In 2014-15 and 2015-16, the period before autumn crop emergence was less dry, with total amounts of rainfall for July and August rainfall of 151 mm in 2014-2015 and 127 mm in 2015-2016. The growing season with the driest summer was the least favorable for pseudothecium maturation, which was delayed by almost two months.

## Discussion

This epidemiological study provides basic information about the interepidemic dynamics of the flax pathogen *M. linicola*, the sexual stage of which was identified in all its forms – pseudothecia, asci, and ascospores – directly on plant host tissues, for the first time in France and in continental Europe. In England, Perryman et al. (2009) trapped airborne *M. linicola* ascospores, but did not observe pseudothecia, raising doubts about the actual presence of the sexual stage on flax residues, and concerning its quantitative significance in particular. Based on our findings and other published evidence, we can now argue that the pathogen survives on flax straw and that sexual reproduction plays a significant role in the epidemiology of the disease. Our epidemiological data suggest that *M. linicola* ascospores produced on flax straw are the major source of primary infection, at least in French conditions. The temporal dynamics of pseudothecium maturity revealed that wind-dispersed ascospores could potentially initiate pasmo epidemics as soon as flax seedlings emerge in the field, in October for winter flax and February-March for spring flax.

Perryman et al. (2009) reported that epidemics began earlier in the growing season when there was much more rainfall in the autumn (1997-98) than growing seasons with drier weather (1998-99, 1999-2000). These findings are consistent with our own, which also suggest a concordance between the earliness of the ascospore peaks (early September 2015 vs. late October 2016) and the earliness of pasmo symptoms (mid-November 2015 vs. mid-December 2016) in field conditions. The dynamics of the *M. linicola* sexual stage on flax residues during the interepidemic period and the pattern of change in pseudothecium maturity over time are consistent with the changes in the concentration of ascospores in the air measured by Perryman et al. (2009) in England and Scotland. Ascospore peaks have been recorded in September and October, the period when the proportion of mature pseudothecia is maximal. Our data, and the few published epidemiological findings available, suggest that ascospores probably play a major role in initiating epidemics, contrary to the prevailing view that pasmo is a seed-borne pathogen (Holmes, 1976).

Our data also suggest that precipitation and humidity are important for pseudothecium maturation, consistent with published findings for other ascomycete pathogens of plants. Temperature is known to affect pseudothecium maturation in ascomycetes, as established, for example, for *L. maculans* and *L. biglobosa* (Toscano-Underwood et al., 2003; Naseri et al., 2009), but this was not demonstrated here because the mean temperatures from early July to late October were similar in the three years considered (16.6°C in 2014-15, 16.5°C in 2015-16 and 17.3 in 2016-17).

The increase in the frequency of pasmo in France in the early 2010s, may, like the increase observed in the UK in the late 1990s (Perryman & Fitt, 2000), reflect an overall increase in the area under flax historically concentrated in the northern part of France, close to those of UK and Belgium (**Figure 5 and 6**). However, it may also reflect an increase in the cultivation of different types of flax crop, resulting in an almost permanent presence of host plants, acting as both a target (living and susceptible tissues) and a source (dead tissues, source of inoculum) of the pathogen. Flax and linseed are sown in the autumn (for winter crops) or the late winter (for spring crops). Winter flax is rarely grown in France (< 1 %), with spring flax the most prevalent form cultivated (85%), followed by winter linseed (12%) and spring linseed (3%) (data CIPALIN; Labalette et al., 2011). In addition to the overall increase in the area under flax crops in France in the three last decades, the diversity of this crop and related practices (but also changes in these practices, e.g. management of straw during the intercropping period) may have account for the size of pasmo epidemics in growing seasons with favorable climatic conditions. First, a resurgence of pasmo in spring flax crops may have occurred over the last two decades in France due to the large amounts of ascospores produced on crop debris as a result of the increase in the area under winter linseed. Pasmo was of little importance on spring linseed before the 1990s in the UK, and Perryman et al. (2009) suggested that the disease would not have become a problem if winter linseed had not provided a source of inoculum. Indeed, straw is poorly degraded and difficult to plough under, since the fibers wrap themselves around disks, wheels and shovels. In the past, the only way to cope with linseed straw was to drop it in windrows after the combine and then burn it directly or harrow or rake it into piles and then burn it. In Canada straw choppers on new combines have been used to effectively chop and spread flax straw, if the straw was dry and relatively short or fiber content was relatively low (Flax Council of Canada, 2015). In UK and France, burning has been practiced for a long time. National regulations still allow to burn flax residues on the ground (e.g. Décret 2015-1769; Legifrance, 2015), but the practice tend to be reduced since the use of straw as biomass energy source is possible. All these changes in management of linseed straw may have played a role in the survival of *M. linicola*.

**Figure 5.**
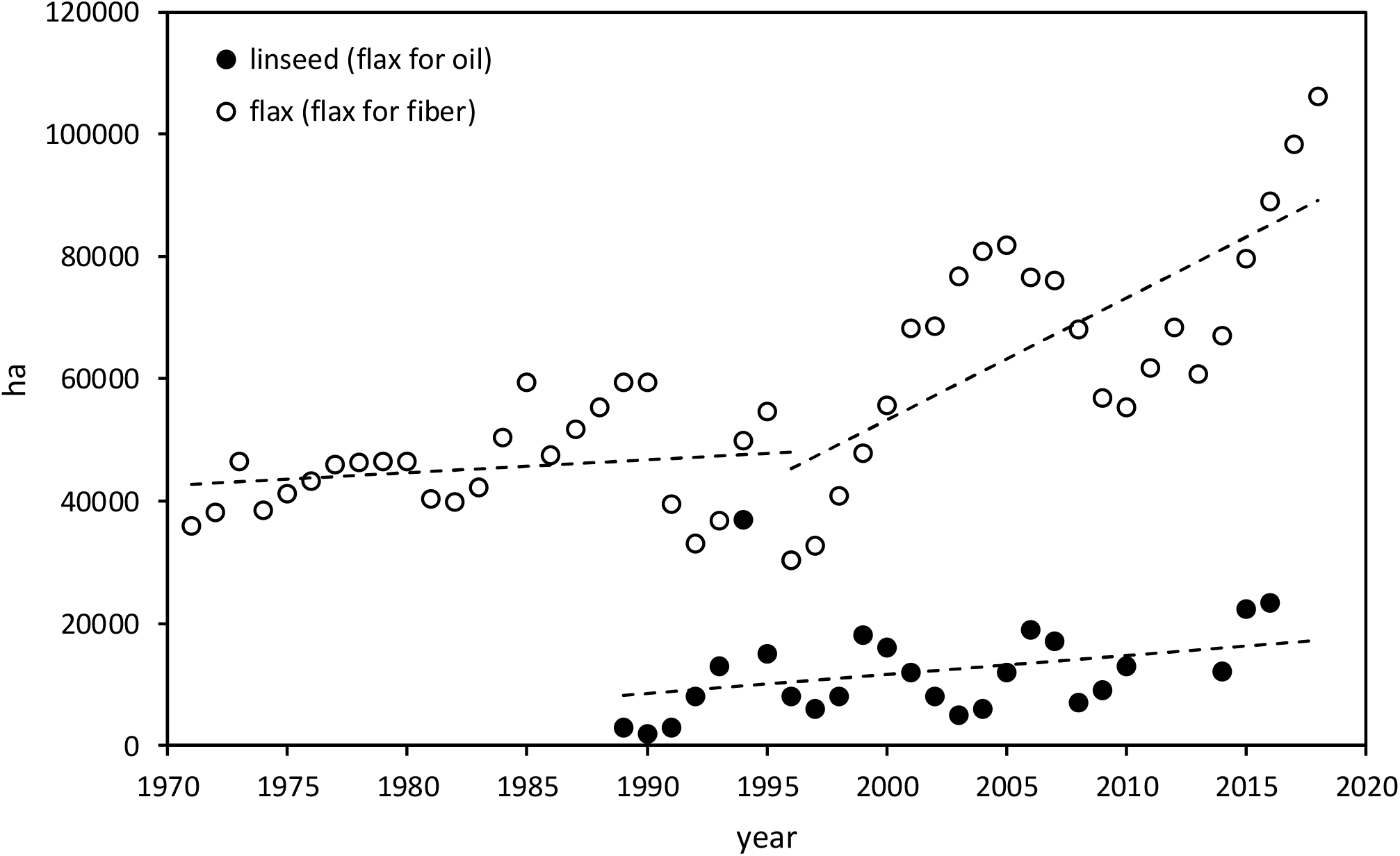
Change in the area under flax/linseed (flax for fiber production and linseed for oil production) in France from 1970 to 2019 (data CIPALIN).

**Figure 6.**
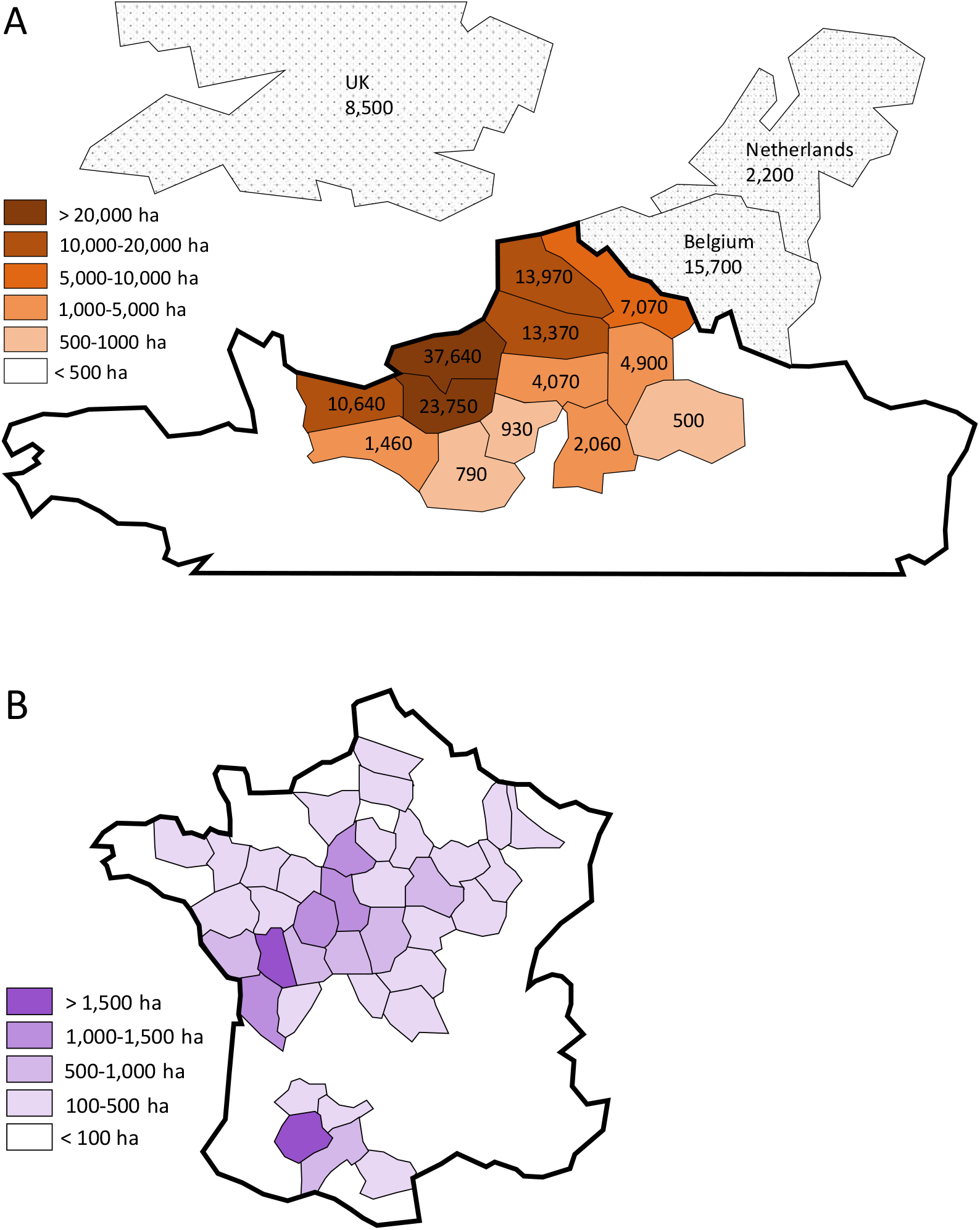
A. Area (ha) under flax for fiber in the main producing French departments and in the main neighbouring producing countries (UK, Belgium and the Netherlands) in 2019 (data CIPALIN and FAOSTAT). B. Area (ha) under linseed in the main producing French departments in 2019 (data CIPALIN).

Second, the increase in pasmo levels in France coincided with a strong increase in the proportion of flax relative to linseed in 1995 (**Figure 5**). It is acknowledged that ploughing or burning crop debris which may carry *M. linicola* inoculum is important to control the disease (Paul et al., 1991), but these practices make sense only for linseed because flax straw is left on the soil for retting (a process in which the action of microorganisms separates the fibers from the other parts of the plant) at the end of summer, sometimes until October. This process may, thus, make a significant contribution to the peak in *M. linicola* ascospore levels just before the emergence of the following crop. Finally, the ‘brown bridge effect’ during the interepidemic period (Kerdraon et al., 2019), boosting the overall amount of inoculum, may be particularly marked as flax and linseed are grown simultaneously in concentrated, close production areas (**Figure 6**), increasing the likelihood of detrimental interactions between sources of inoculum and target plant populations.

Third, resistance tests performed in controlled conditions within the framework of the ‘SeptoLIN’ project have highlighted significant differences in pasmo sensitivity between the main linseed (n = 7) and flax (n = 15) cultivars currently grown in France, the latter presenting higher levels of disease (Penaud et al., 2017). This significant difference (p < 0.05), assuming that it was stable over time, could also have contributed marginally to the increase in the frequency of pasmo.

This study illustrates how a particular part of the disease cycle, in this case, the sexual phase, can become crucial due to changes in agronomic practices and the processing of plant products. The epidemiological consequences of such changes to the production and processing system must be taken into account at larger spatiotemporal scales, to improve crop protection strategies. Several fungal diseases of agronomic importance caused by ascomycetes have a direct impact during the epidemic phase (yield reduction due to effects on plant growth) and an indirect impact during the interepidemic phase (increase in inoculum production and disease pressure at early stages in the development of the following crop). In the case of flax, there may be a double direct effect on both the period of cultivation and the interepidemic period, because sexual reproduction takes place on the valuable part of the plant – the stem, from which fibers are extracted – during retting. *Verticillium* wilt, another important flax disease caused by the soilborne fungus *Verticillium dahliae*, is known to damage flax fibers and to cause significant yield losses. This fungal pathogen reaches the fiber during retting, leading to the embedding of numerous microsclerotia within the bast fiber bundle of the stem, which becomes brittle and fragile (Blum et al., 2018). Asexual infections of the stems with *M. linicola* do less damage to the fibers than *V. dahliae*, and no specific impact of sexual reproduction (i.e. the formation of pseudothecia within the fibers) has yet been established. However, the issue of possible damage to the fibers due to pasmo is of interest. In particular, it would be useful to estimate the biological, chemical and physical impacts of *M. linicola* on the retting process in the field, fiber quality and the production of inoculum for the infection of flax plants in the following cropping period.

We show here that the sexual stage of *M. linicola* is of great epidemiological importance in pasmo. This importance may have increased over the last two decades. Our findings suggest that the changes observed in flax production may have resulted in *M. linicola* injuring the plant in two ways, depending on the production conditions (Savary et al., 2000; 2012) and assessments of the yield losses caused by pasmo (Perryman & Fitt, 2000). Approaches taking this damage into account pave the way for the maintenance of scutched flax fiber quality, and, more generally, the sustainability of the French linen/linseed sector, through the control of pasmo without the need for an increase in fungicide use.

---

**Supplementary Figure 1.**
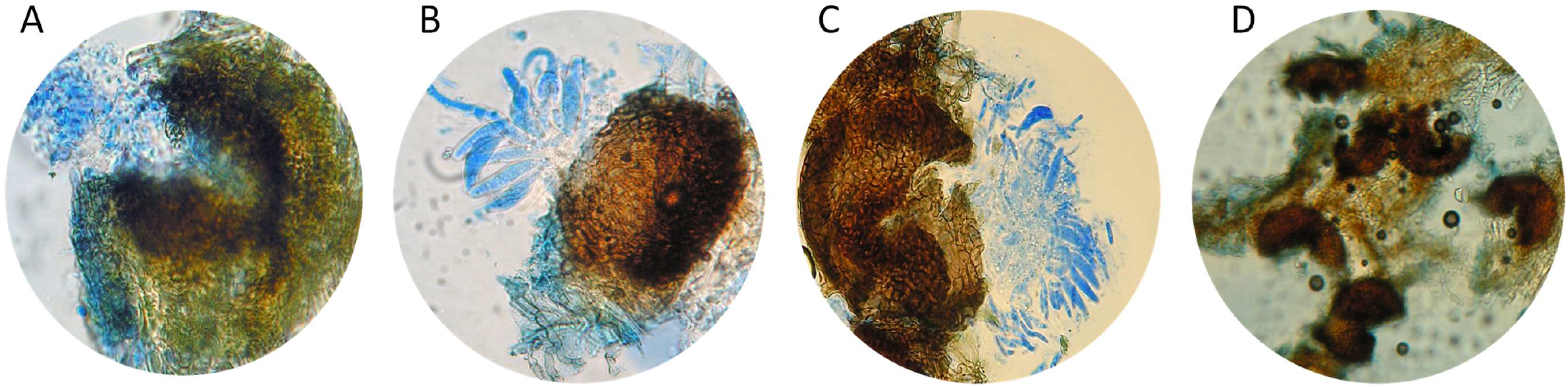
Visual categories for recognition and classification of the maturation of each pseudothecium of *Mycosphaerella linicola* based on the stage of development of asci and ascospores (i.e. degree of maturity): A. Immature; B. Maturing; C. Mature; D. Empty.

**Supplementary Figure 2.**
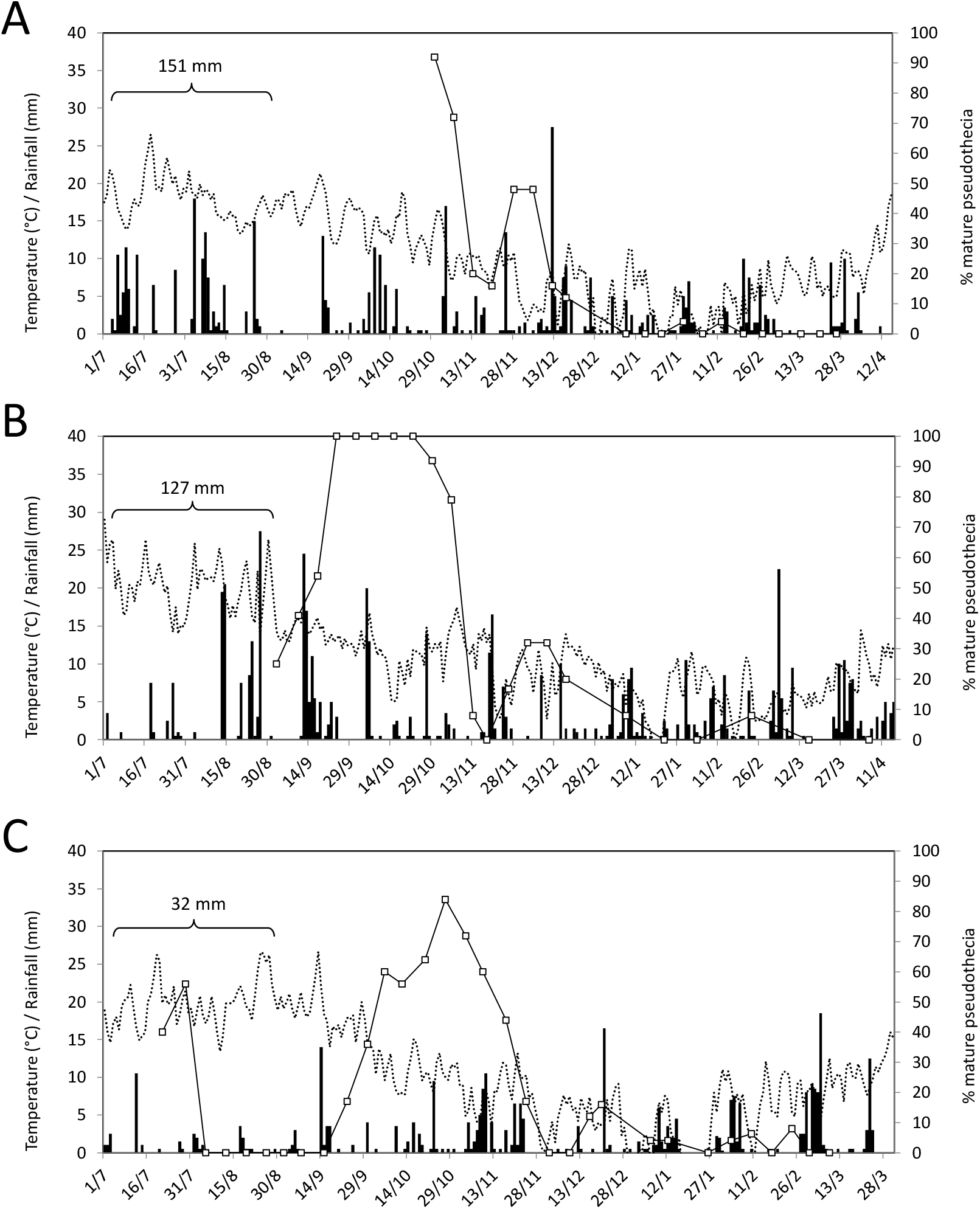
Mean daily air temperature (dotted line, °C) and daily rainfall (histogram, mm) at Grignon from July to April in the 2014 (A), 2015 (B) and 2016 (C) growing seasons, relative to the change in the percentage of mature *Mycosphaerella linicola* pseudothecia on flax straws over time (black line). Cumulative rainfall in the summer, which is thought to have a strong impact on the early dynamics of the sexual stage of *M. linicola* on flax straw, is indicated.

## References

Adee, E.A., Pfender, W.F. (1989). The effect of primary inoculum level of Pyrenophora tritici-repentis on tan spot epidemic development in wheat. Phytopathology 79: 873–877.

Blum, A., Bressan, M., Zahid, A., Trinsoutrot-Gattin, I., Driouich, A., Laval, K. (2018). Verticillium wilt on fiber flax: symptoms and pathogen development in planta. Plant Disease 102: 2421–2429.

Brentzel, W.E. (1926). The pasmo disease of flax. Journal of Agricultural Research 32: 25–37.

Christenson, I.J. (1952). Dissemination of Septoria linicola in relation to epidemics of pasmo. Proceedings of the 33’d Flax lnstitute of the United States. USA.

Colhoun, J, Muskett, A.E. (1943). “Pasmo” disease of flax. Nature 151: 223–224.

Ferguson, M.W., Lay, C.L., Evenson, P.D. (1987). Effect of pasmo disease on flower production and yield components of flax. Phytopathology 77: 805–808.

Fitt, B.D.L., Jouan, B., Sultana, C., Paul, V.H., Bauers, F. (1991). Occurrence and significance of fungal diseases on linseed and fibre flax in England, France, and Germany. Aspects of Applied Biology 28: 59–64.

Gillis, E.O. (2009). Investigating partial resistance and host-pathogen interactions in the flax – Septoria linicola pathosystem. MSc thesis, University of Manitoba, Canada.

Flax Council of Canada (2015). Chapter 12. Flax straw and fibre – Past and present uses. In: Growing flax – Production, management and diagnostic guide. https://flaxcouncil.ca/wp-content/uploads/2015/02/FCOC-growers-guide-Chapter-12-Flax-Straw-and-Fibre.pdf

Grant, L. (2008). Effect of pasmo on flax in Manitoba and inference of the sexual state of the fungus by molecular polymorphism. PhD thesis, University of Manitoba, Canada.

Holmes, S.J. (1976). Pasmo disease of linseed in Scotland. Plant Pathology 25: 61.

Kerdraon, L., Laval, V., Suffert, F. (2019). Microbiomes and pathogen survival in crop residues, an ecotone between plant and soil. Phytobiome Journal 3: 246–255.

Kruger, E. (1941). Untersuchungen über zwei der bedeutendsten Leinparasiten - Colletotrichum lini Manns et Bolley und Septoria linicola (Speg.) Gar. (Sphaerella linicola Wr.). Arb. Biol. Reichsanst. Ld-u. Forstw. 23: 163–188.

Labalette, F., Landé, N., Wagner, D., Roux-Duparque, M., Saillet, E. (2011). La filière lin oléagineux française : panorama et perspectives. OCL 18: 113–122.

Legifrance (2015). Décret n° 2015-1769 du 24 décembre 2015 relatif aux bonnes conditions agricoles et environnementales des terres. Texte n°97. Journal officiel de la République française 0300:24184

Loughnane, J.B., McKay, R., Lafferty, H.A. (1946). Observations on the pasmo disease of flax and on the causal organism Mycosphaerella linorum Wollenweber. The Scientific Proceedings of the Royal Dublin Society 24: 89–98.

Morais, D., Sache, I., Suffert, F., Laval, V. (2016). Is onset of Septoria tritici blotch epidemics related to local availability of ascospores? Plant Pathology 65: 250–260.

Naseri, B., Davidson, J.A., Scott, E.S. (2009). Maturation of pseudothecia and discharge of ascospores of Leptosphaeria maculans on oilseed rape stubble. European Journal of Plant Pathology 125: 523–531.

Paul, V.H., Sultana, C., Jouan, B., Fitt, B.D.L. (1991). Strategies for control of diseases on linseed and fibre flax in Germany, France and England. Aspects of Applied Biology 28: 65–70.

Penaud, A., Paumier, D., Bamme, B., Petiteau, A., Heritier, E., Suffert, F., Valade, R. (2017) Epidemiology of pasmo and Septoria linicola resistance in French flax cultivars (abstract). Proceedings of the 12th EFPP-10th SFP Conference, May 29 to June 2, 2017, Malo-les-Bains, France.

Perryman, S., Fitt. B.D.L. (2000). Effects of diseases on the yield of winter linseed. Aspects of Applied Biology 56: 211–218.

Perryman, S., Gladders, P., Fitt, B.D.L. (2009). Autumn sowing increases severity of pasmo (Mycosphaerella linicola) on linseed in the UK. Annals of Applied Biology 154: 19–32.

Poisson, B. (1997). Etudes relatives à la maturation des périthèces de Leptosphaeria maculans sur les pailles de colza d’hiver nécrosées au collet. 5ème Conférence sur les Maladies des Plantes, Tours, France. 1:345–352.

Rost, H. (1937). The pasmo disease of flax in Europe caused by Septoria linicola (Speg.) Garassini. Review of Applied Mycology 16: 676–677.

Sanderson, F.R. (1963). An ecological study of pasmo disease (Mycosphaerella linorum) on linseed in Canterbury and Otago. New Zealand Journal of Agricultural Research 6: 432–439.

Sivaneson, A., Holliday, P. (1981). Mycosphaerella linicola. Descriptions of fungi and bacteria. Commonwealth Mycological Institute, UK.

Sackston, W.E. (1949). Studies on the pasmo disease of flax. PhD thesis, University of Minnesota, St. Paul, USA.

Sackston, W.E., Carson, R.B. (1951). Effect of pasmo disease of flax on the yield and quality of linseed oil. Canadian Journal of Botany 29: 339–351.

Sackston, W.E. (1970). A possible mechanism of dispersal of Septoria spores. Canadian Journal of Plant Science 50: 155–157.

Savary, S., Ficke, A., Aubertot, J.N., Hollier, C. (2012). Crop losses due to diseases and their implications for global food production losses and food security. Food Security 4: 519–537.

Savary, S., Willocquet, L., Elazegui, F.A., Castilla, N.P., Teng, P.S. (2000). Rice pest constraints in tropical Asia: quantification of yield losses due to rice pests in a range of production situations. Plant Disease 84: 357–369.

Sivaneson, A., Holliday, P. (1981). Mycosphaerella linicola. In: Descriptions of fungi and bacteria. Commonwealth Mycological Institute, UK.

Spegazzini, C. (1911). Phtyctena ? linicola Speg. (n.f.). Mycetes Argentinenses. Annales del Museo Nacional de Buenos Aires, Serie III 13: 389–390.

Suffert, F., Sache, I. (2011). Relative importance of different types of inoculum to the establishment of Mycosphaerella graminicola in wheat crops in north-west Europe. Plant Pathology 60: 878–889.

Toscano-Underwood, C., Huang, Y.J., Fitt, B.D.L., Hall, A.M. (2003). Effects of temperature on maturation of pseudothecia of Leptosphaeria maculans and L. biglobosa on oilseed rape stem debris. Plant Pathology 52: 726–736.

Vakhrusheva, T.A. (1986). Ascous stage of pasmo affecting flax plants (in Russian). Zashchita rastenii 29.

Wollenweber, H.W. (1938). “Sphaerelle linicola” N. sp. Die Ursache der Amerikanischer Leinpest. (Pasmo - oder “Septoría” - Krankheit.). Revista de Botanica del Instituto Miguel Lillo, Lilloa 2: 483–495.

